# Understanding binary classifier model structure based on Shapley feature interaction patterns

**DOI:** 10.1101/2021.03.29.437591

**Authors:** Boyang Zhao

## Abstract

There is increasing emphasis on the interpretability of machine learning models, including in understanding biological systems. The well-known Shapley value framework based on game theory works in principle with any models to attribute feature importance. While feature interactions are critical to understand and can be interpreted within this framework, much attention is paid in practice on global feature importance and general trends of interactions. The inter-relationships between underlying model structure and Shapley value and its decomposition is less clear. Here we use binary classifiers to systematically examine how logical and additive interactions affect marginal contributions. These decomposed main and interaction effects are reflected in resulting Shapley dependence plots. The directionality of inequalities or logical/additive operators influence independently the main and marginal/interaction effects. Lastly, we show that these principles are applicable for models with noise.

## 1 Introduction

Model explainability is an important part of machine learning and critical in understanding factors that are driving a model’s decision. This includes using models to understand genomic drivers of cellular phenotypes. A myriad of techniques have been developed, including model-agnostic approaches such as SHAP^1^, which is based on Shapley values^2^. However, often the emphasis is placed on a ranked list of global feature importance with less attention paid to feature interactions. At best feature interactions are at the level of understanding a feature’s main effect and basic trends of feature interaction. The question remains in how different model structures relate to for example the often used Shapley values, its marginal contributions, and dependence plots.

While several approaches exist for assessing feature interactions such as H-statistics^3^, partial dependence plot-based variable importance^4^, variable interaction networks^5^, etc, we focus primarily on Shapley/SHAP interactions. This is because of their wide spread usage and they are based on a strong foundations of game theory and are model agnostic. Here, we systemically build binary classifiers with different feature interactions, and observe its impact on Shapley values and contributions.

## 2 Feature contribution with Shapley values

Shapley values was developed as an approach in cooperative game theory to estimate the contribution of each player in a coalitional game consisted of multiple players. In the context of machine learning, an individual player corresponds to a feature in a model. Let ***x*** ∈ ℝ^*M*^ be the *M* features in the model.

The Shapley value for the *i*th feature is given by

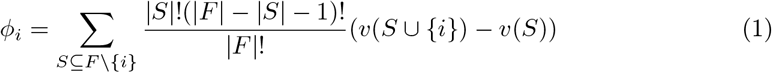

where *F* is the set of all *M* features and *S* is a subset of the features (*S* ⊆ *F*). The function *v*(*S*) is a characteristic function that returns the expected payoffs of a given coalition (i.e. model evaluation based on a subset of features). For each given feature subset *S*, we calculate the payoffs of coalitions of *S* with and without the ith feature. The difference of the two represents the marginal contribution of the ith feature given *S*, of which we weight over all permutations in which the coalition of *S* can be formed. This is repeated to sum over all feature subsets not containing the *i*th feature. The resulting Shapley value describes the contribution (or impact) of the *i*th feature of a given model.

Lundberg and Lee^1^ introduced SHAP values by applying Shapley values to machine learning models and defined the characteristic function *v* as a conditional expectation function,

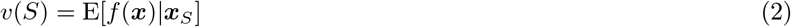

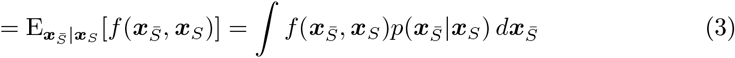

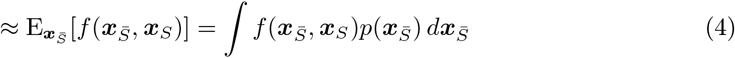

Here the *v*(*S*) is the expectation of the model given coalition of subset features *S*, which is calculated by marginalizing out the other features 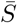. We simplify Equation 3 to 4 by assuming feature independence^1^. Our discussions below use the approximation Equation 4 when referring to Shapley values. For computational efficiency, several approximation methods (e.g. Kernel SHAP^1^, Tree SHAP^7^) have also been introduced for estimating SHAP values.

The Shapley/SHAP values defined thus far relate to the total effect of a single feature. The marginal contribution of the feature in a particular feature set is described in *v*(*S*∪{*i*}) – *v*(*S*). In addition to the total and marginal effects, the effects can also be decomposed into main effects and interaction effects. The SHAP interaction values is based on the Shapley interaction index^8;9^, and is defined as^7^,

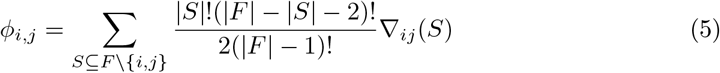

where *i* ≠ *j* and,

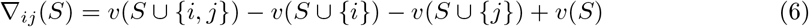

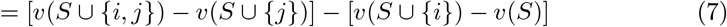

The intuition is that the interaction effect is the difference between the marginal effect of *x_i_* with and without *x_j_*. Equation 7 can be rearranged so the marginal effect is of *x_j_*. The total interaction is split equally between *ϕ_i,j_* and *ϕ_i,j_*.

The main effect can be derived as the difference between the total effect *ϕ_i_* and the interaction effects *ϕ_ij_*,

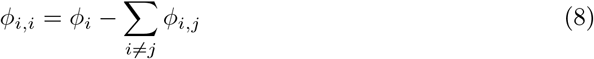

## 3 Binary classifier model

We will use a two feature binary classifier model (***x*** ∈ ℝ^2^), with varying interaction types, to examine their effects.

For *i* = 1, there are two feature subsets consisting of 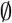 and {2}. The Shapley value for *x*_1_ is,

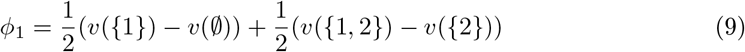

For *i* = 2, the feature subsets consist of 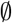 and {1}, with the Shapley value as,

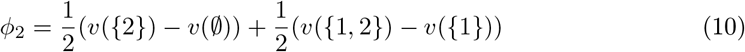

We consider *x* ~ *U*(0,1). Given this, the characteristic functions per approximation in Equation 4 are,

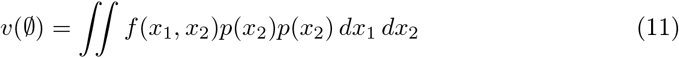

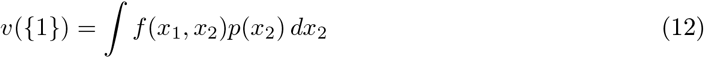

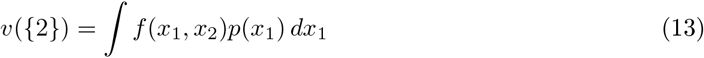

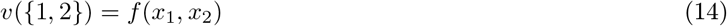

When we examine the characteristic functions of each feature subsets, we can conclude already that 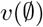 is a constant; *v*({*i*}) is related to the main effects of *x_i_* as *x_j_* is integrated out^2^, and we will see later this is indeed the case when examining main vs interaction effects; and *v*({1, 2}) is just *f*(***x***). As such, when we decompose *ϕ_i_* into the marginal effect components 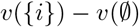 and *v*({1, 2}) – *v*({*j*}), the first component is the marginal effect of *x_i_* relative to a null feature set and the second component is the marginal effects of *x_i_* relative to a *x_j_* feature set.

We can also directly decompose *ϕ_i_* into its main and interaction effects. The interaction effect between *x*_1_ and *x*_2_ (Equation 5) is,

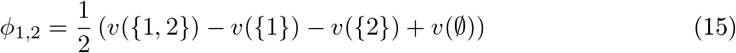

With *ϕ*_1,2_ and *ϕ_i_*, the main effects (Equation 8) are,

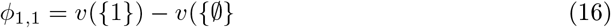

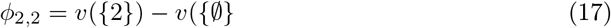

By this formulation, the main effect of *x_i_* is in fact the same as the marginal component 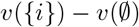 discussed earlier. Taken together, these mean that changes to the main effects of feature *x_i_* is reflected in *v*({*i*}) and thus the main effect/marginal effect term 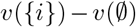; while changes to the feature interactions is reflected in *v*({1,2}) and consequently the interaction effect *ϕ*_1,2_ and the marginal effect term *v*({1,2}) – *v*({*j*}).

## 4 Feature interactions

We can now define *f*(***x***) with different *x*_1_ and *x*_2_ interactions. We will first examine logical relations between *x*_1_ and *x*_2_, starting with an AND relation, and examine how changing this to an OR relation and switching of the inequality signs affect our characteristic functions and resulting dependence plots.

### 4.1 AND operation

Suppose we start with our two-feature model with the following *f*(***x***),

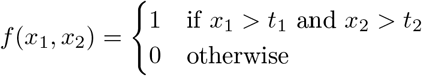

The characteristic functions are,

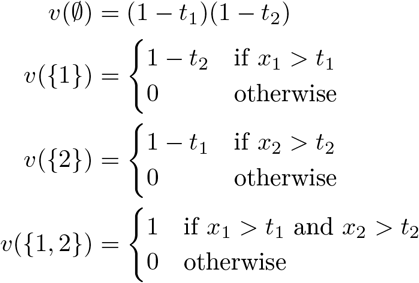

The Shapley value for each feature can be decomposed into two components: 1) the marginal contribution of feature to a null-set coalition, 2) the marginal contribution of feature to a coalition consisted of the other feature. For *ϕ*_1_, this corresponds to 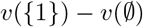 and *v*({1,2}) – *v*({2}), respectively.

Combining these two components, as described in Equation 9 and 10, the Shapley values for the two features are as follows,

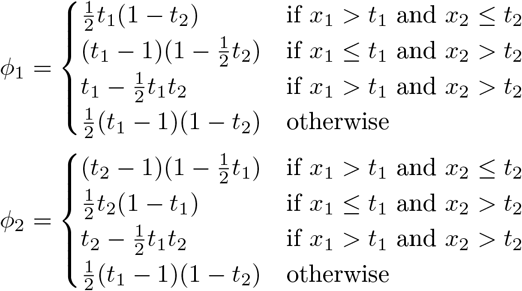

As an example below, we will use, without loss of generality, *t*_1_ = 0.8 and *t*_2_ = 0.4. This corresponds to an upper right region in the feature space where *f*(***x***) is equal to 1 (Figure 1B). The characteristic function of individual feature subsets concur with our discussions in Section 3, viz. 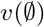 is a constant, *v*({*i*}) corresponds to the individual effects of *x_i_*, and *v*({1, 2}) is *f*(***x***) (Figure 1C).

**Figure 1:**
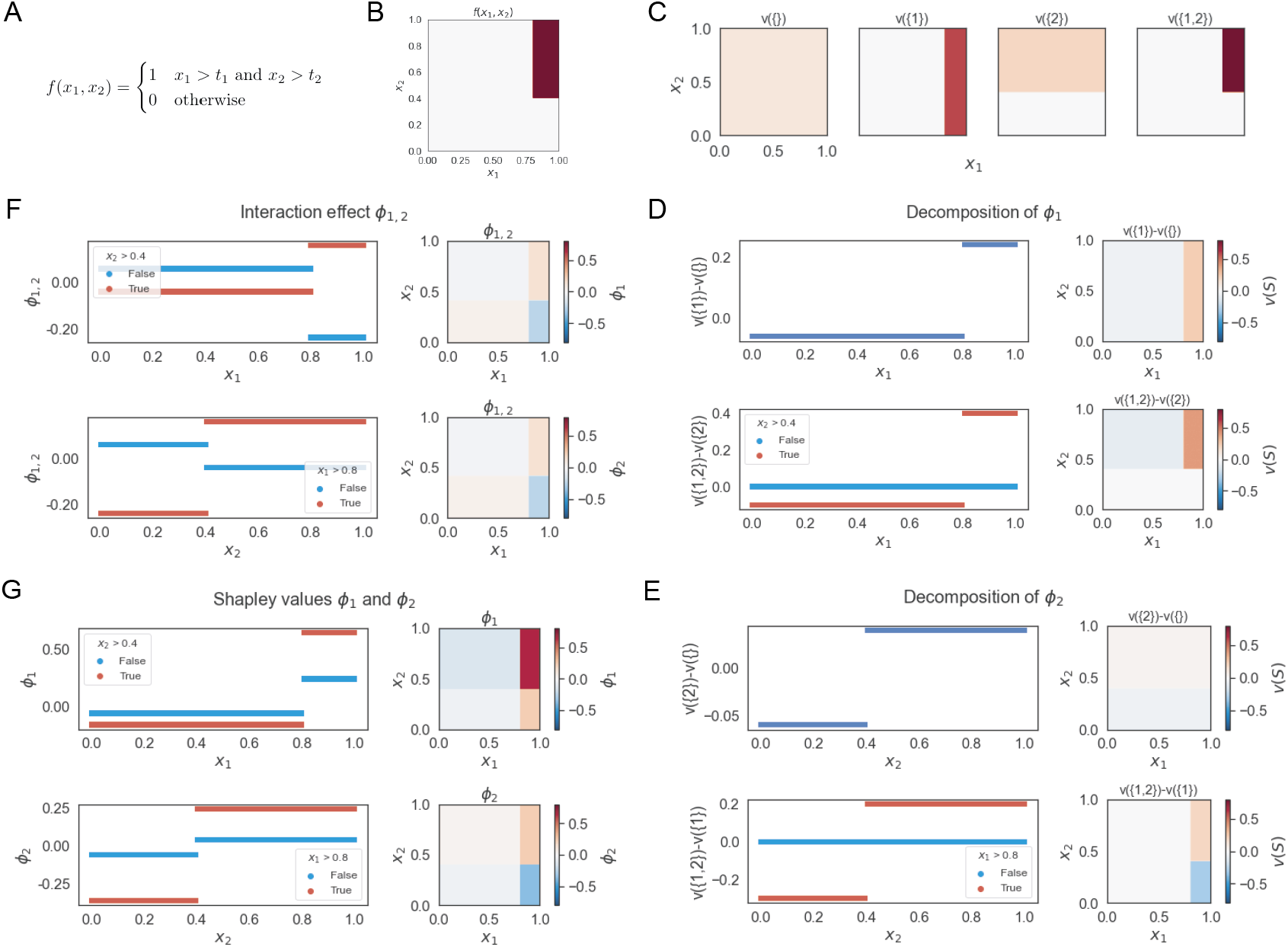
Summary of Shapley values for two-feature binary classifier with AND feature relation. (A) Classifier *f*(***x***) takes the value 1 when both *x*_1_ and *x*_2_ are above their thresholds. *x* ~ *U*(0,1). (B) Heatmap of *f*(***x***) where dark red region indicates *f*(***x***) = 1. (C) Heatmap of each characteristic functions. (D-F) Decomposition of Shapley values into the marginal contributions 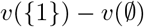 and *v*({1, 2}) – *v*({2}), for *x*_1_ (D) and *x*_2_ (E), and interaction effects (F). (G) Shapley values for *x*_1_ and *x*_2_ as dependence plots and heatmaps.

When we examine the decomposition of *ϕ*_1_, we expectedly observe that the marginal effect 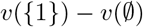 describes the main effects of *x*_1_. The marginal effect *v*({1,2}) – *v*({2}) describes the additional contribution after having the feature *x*_2_ be already added to the coalition. With an ADD relation, this means if *x*_2_ ≤ *t*_2_, there is no value of *x*_1_ that will change *f*(*x*) to equal 1 as AND requires also *x*_2_ > *t*_2_; consequently the marginal contribution of *x*_1_ is zero. On the other hand, when *x*_2_ > *t*_2_, values of *x*_1_ above threshold *t*_1_ exhibits a positive marginal effect, while below threshold a negative marginal effect (Figure 1D). The same behavior is observed for *x*_2_ (Figure 1E).

Taken together, we can easily see both the main effects of *x_i_* and the marginal effects of *x_i_* given *x_j_* reflected in the Shapley values *ϕ_i_* (Figure 1G). With an ADD feature interaction, when *x_j_* > *t_j_* (this is ’in-region’ where other feature matters in an ADD relation), *x_i_* will appear to *magnify* the resulting Shapley values in dependence plots for *ϕ_i_* as compared to *x_j_* ≤ *t_j_*. Furthermore, when both features are above or below their thresholds, they have positive Shapley interaction values; and when one is above and the other is below, they have negative interaction values (Figure 1F).

### 4.2 OR operation

Now let us see what is the effect of changing AND to an OR relation. Here *f*(***x***) can be defined as,

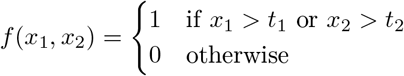

The characteristic functions are,

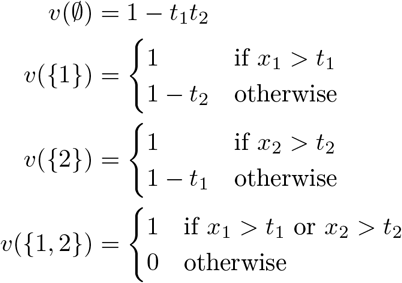

With this, we can derive our Shapley values,

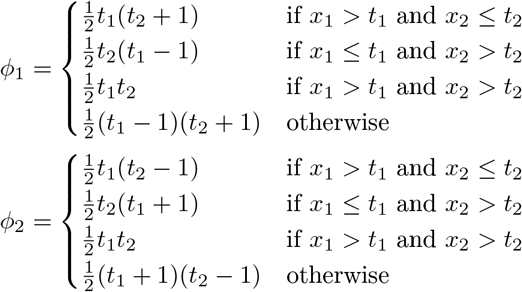

Per discussions in Section 3, we note the main/marginal effect 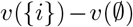 is independent of the *x*_1_ and *x*_2_ interactions, whereas the marginal effect *v*({1, 2}) – *v*({*i*}) is dependent on the interaction. As a result, we expectedly observe that the main effects of *x*_1_ remain unchanged, but the marginal effects of *x*_1_ given *x_j_* (and interaction effect) to be switched when we change the feature relations from AND to OR. Unlike in the AND relation, if *x*_2_ > *t*_2_, *f*(***x***) is equal to 1 independent of the values of *x*_1_ and therefore the marginal contribution of *x*_1_ given *x*_2_ > *t*_2_ is zero. Conversely, if *x*_2_ ≤ *t*_2_, then *f*(***x***) is fully dependent on *x*_1_ (Figure 2).

**Figure 2:**
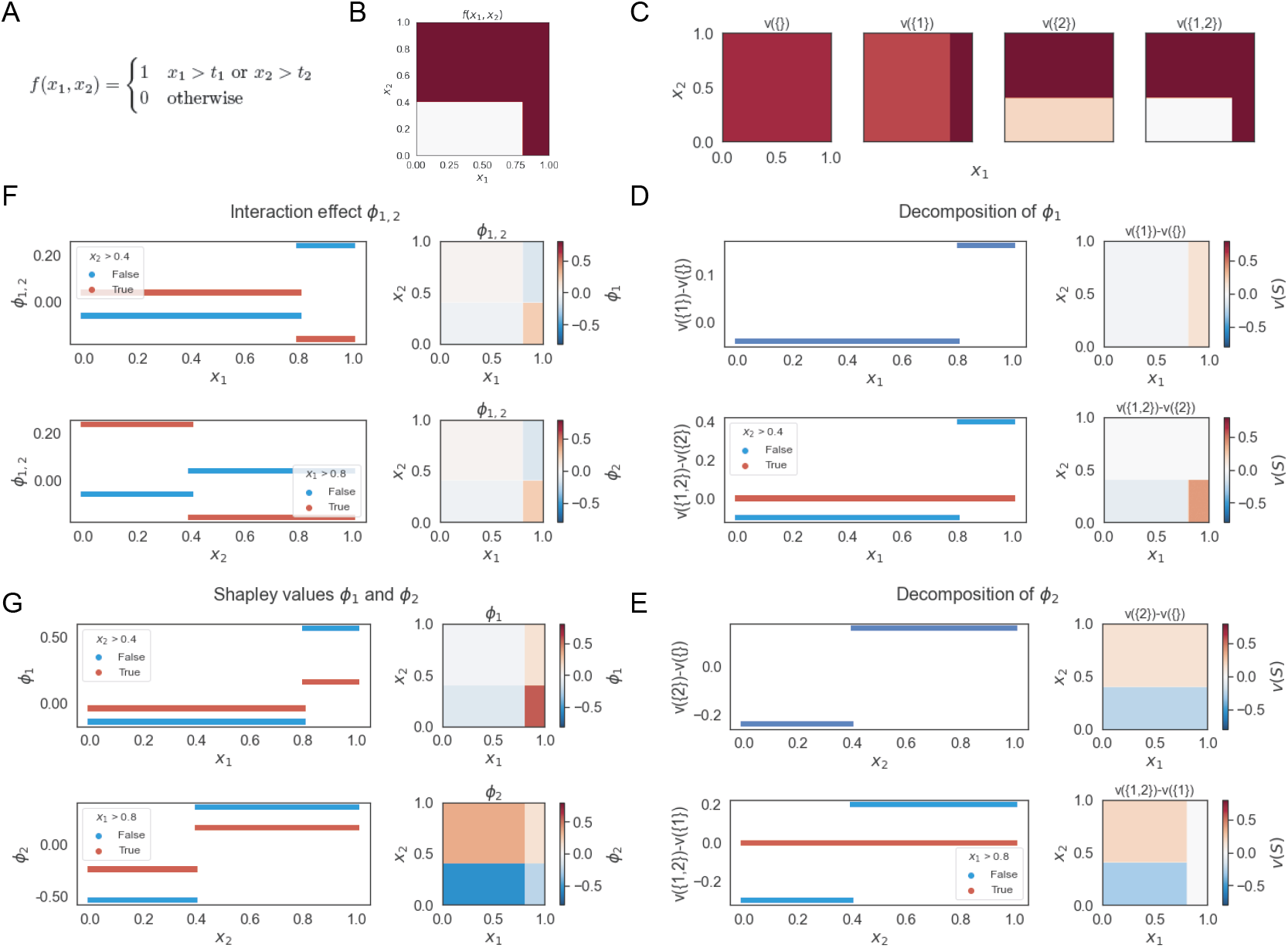
Summary of Shapley values for two-feature binary classifier with OR feature relation. (A) Classifier *f*(***x***) takes the value 1 when either *x*_1_ or *x*_2_ are above their thresholds. *x* ~ *U*(0,1). (B) Heatmap of *f*(***x***) where dark red region indicates *f*(***x***) = 1. (C) Heatmap of each characteristic functions. (D-F) Decomposition of Shapley values into the marginal contributions 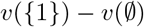 and *v*({1, 2}) – *v*({2}), for *x*_1_ (D) and *x*_2_ (E), and interaction effects (F). (G) Shapley values for *x*_1_ and *x*_2_ as dependence plots and heatmaps.

Thus, with an OR relation, when *x_j_* ≤ *t_j_*, it is ’in-region’ where *x*_1_ matters and *x*_1_ will appear to *magnify* the resulting Shapley values in dependence plots for *ϕ_i_*, as compared to *x_j_* > *t_j_* (Figure 2G). Furthermore, when both features are above or below their thresholds, they have negative Shapley interaction values; and when one is above and the other is below, they have positive interaction values (Figure 2F).

### 4.3 Inequality sign switch

If the main effects are reflected in the marginal contribution of 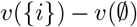, this also means this component is directly affected when we change the direction of inequality. Take for example the function,

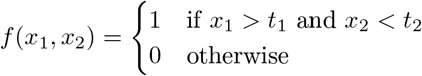

The main effects of *x_i_* can be readily observed in the individual *v*({*i*}) characteristic functions and the Shapley dependence plots (Figure 3). In fact, we can also easily identify that *x*_1_ is associated with a *greater than* inequality and *x*_2_ is associated with a *less than* inequality based on their respective main effects.

**Figure 3:**
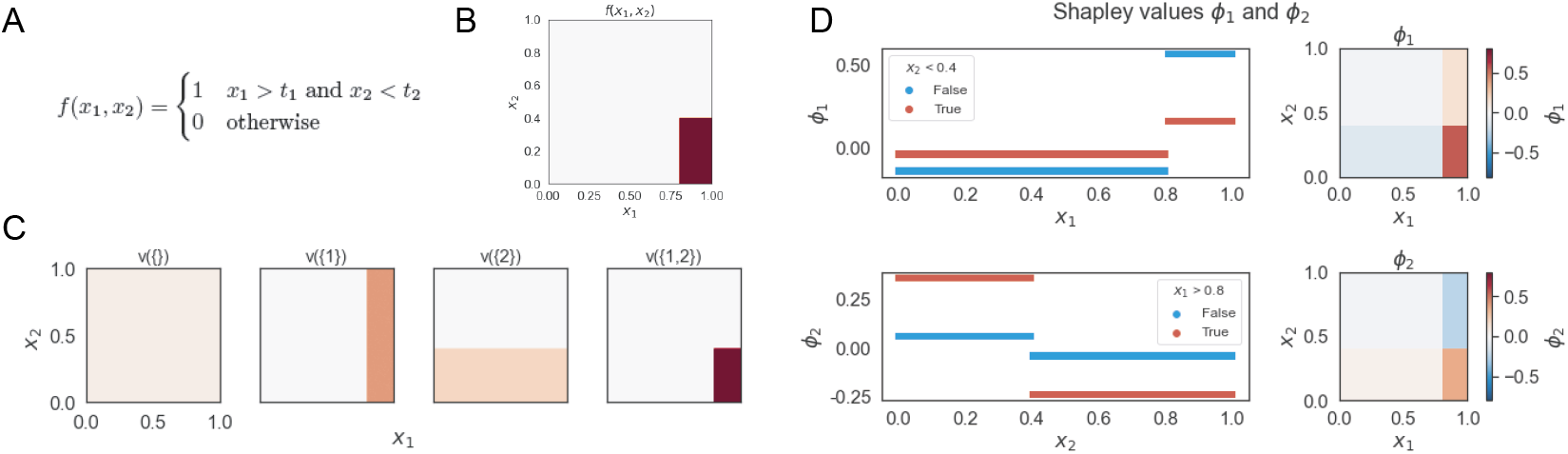
Summary of Shapley values for two-feature binary classifier with AND feature relation; similar to Figure 1, but with inequality direction switched for *x*_2_. (A) Classifier where *f*(***x***) takes the value 1 when both *x*_1_ and *x*_2_ are above and below their thresholds, respectively. *x* ~ *U*(0,1). (B) Heatmap of *f*(***x***) where dark red region indicates *f*(***x***) = 1. (C) Heatmap of each characteristic functions. (D) Shapley values for *x*_1_ and *x*_2_ as dependence plots and heatmaps.

Furthermore, we note from the dependence plots that when *x*_1_ > *t*_1_ or *x*_2_ < *t*_2_, with an AND relation, they are ’in-region’ and thus changes to correspondingly *x*_2_ or *x*_1_ *magnifies* the Shapley values *ϕ*_2_ or *ϕ*_1_, respectively (Figure 3D).

### 4.4 Additivity

Aside from logical operations, we can also construct feature interactions based on arithmetic operations. Suppose we have the following additive feature interaction,

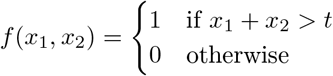

The characteristic functions are,

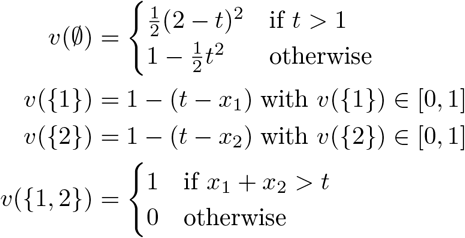

In contrast to the logical operations, in the functions with arithmetic operations, the Shapley dependence plots is not a step function (Figure 4) but instead exhibit linear relationships between *ϕ_i_* and *x_i_* in defined regions. Our conclusions regarding main vs. interaction effects as discussed in Section 3 are still applicable. Namely the main effects of *x_i_* are reflected in 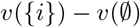. The marginal contribution *v*({1, 2}) – *v*({*i*}) and interaction effect describes the additive interaction between the two features. The region for which *x*_1_ + *x*_2_ > *t* results in higher Shapley values. Moreover, in this region, a lower value of xj results in a higher Shapley value for *ϕ_i_* because *x_i_* now needs to contribute more to get past the threshold *t*. There are also some effects of whether *t* > 1 or *t* < 1, but aside from when the linearity appears, the resulting dependence plots are consistent with the earlier observations.

**Figure 4:**
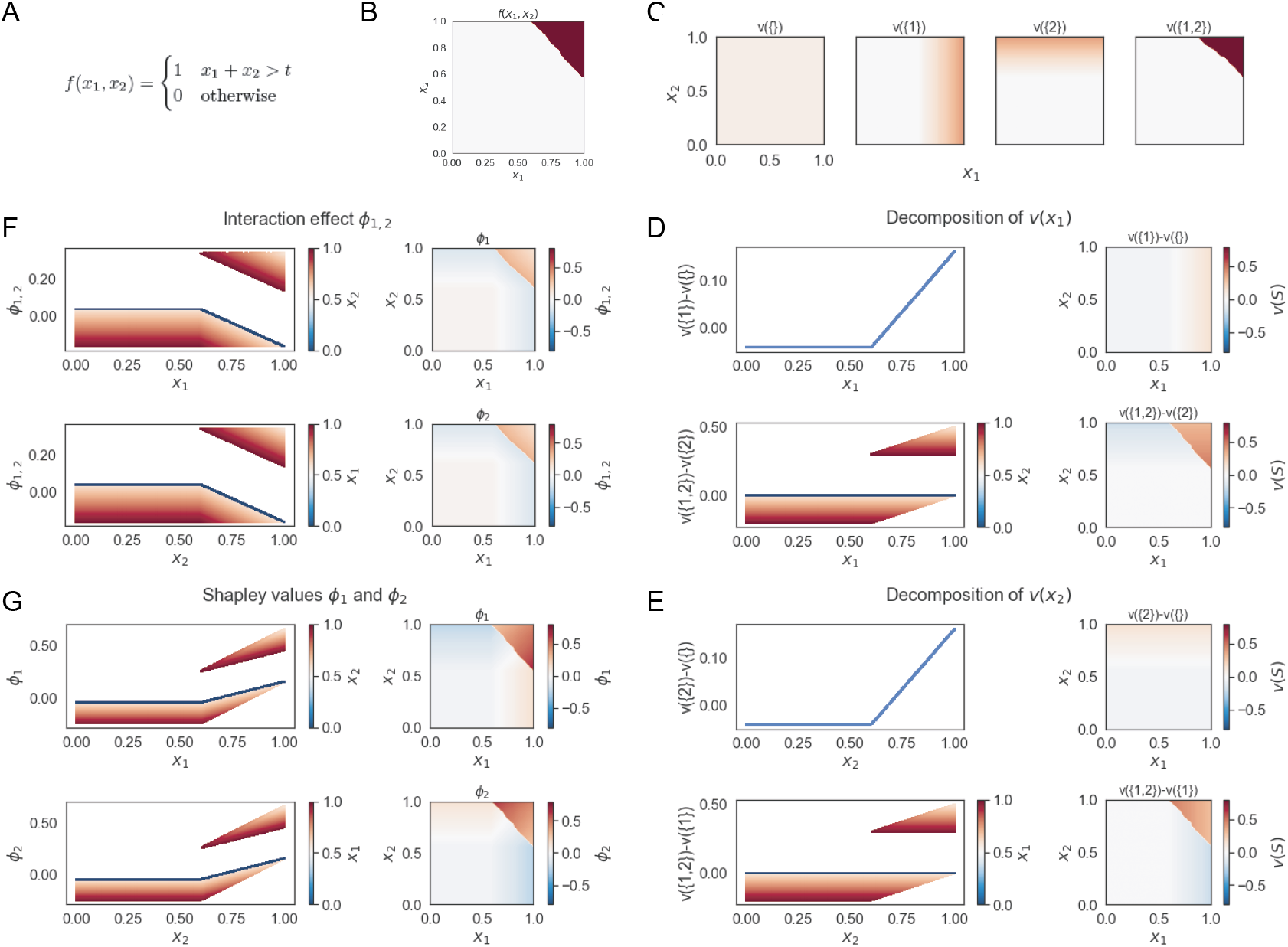
Summary of Shapley values for two-feature binary classifier with additive feature relation. (A) Classifier where *f*(***x***) takes the value 1 when the sum *x*_1_ and *x*_2_ are above threshold t. *x* ~ *U*(0,1). (B) Heatmap of *f*(***x***) where dark red region indicates *f*(***x***) = 1. (C) Heatmap of each characteristic functions. (D-F) Decomposition of Shapley values into the marginal contributions of 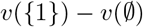 and *v*({1, 2}) – *v*({2}), for *x*_1_ (D) and *x*_2_ (E), and interaction effects (F). (G) Shapley values for *x*_1_ and *x*_2_ as dependence plots and heatmaps.

Several other effects are worth mentioning. As previously discussed in Section 4.3, switching of the inequality sign correspondingly affects the main effects of each feature. The marginal effects of *x_i_* given *x_j_* and the interaction effects remain additive in nature. In all, the logics in the interrelationships amongst the different contribution components and Shapley values are the same as discussed.

## 5 Models with noise

Thus far, we have used noiseless models to gain insights into model structure based on characteristic functions, marginal effects, and Shapley values. These interconnections are retained when we consider models with noise. For this we set a background noise such as *f*(***x***) ~ Bern(*p*). For approximations, we build a tree ensemble random forest model with feature contributions calculated using SHAP^3^.

We can distinctly observe that the underlying model for Figure 5A exhibits a logical AND feature interaction whereas that for Figure 5B exhibits additive interaction. The spreads of SHAP values in the dependence are consistent with our earlier discussions of these models.

**Figure 5:**
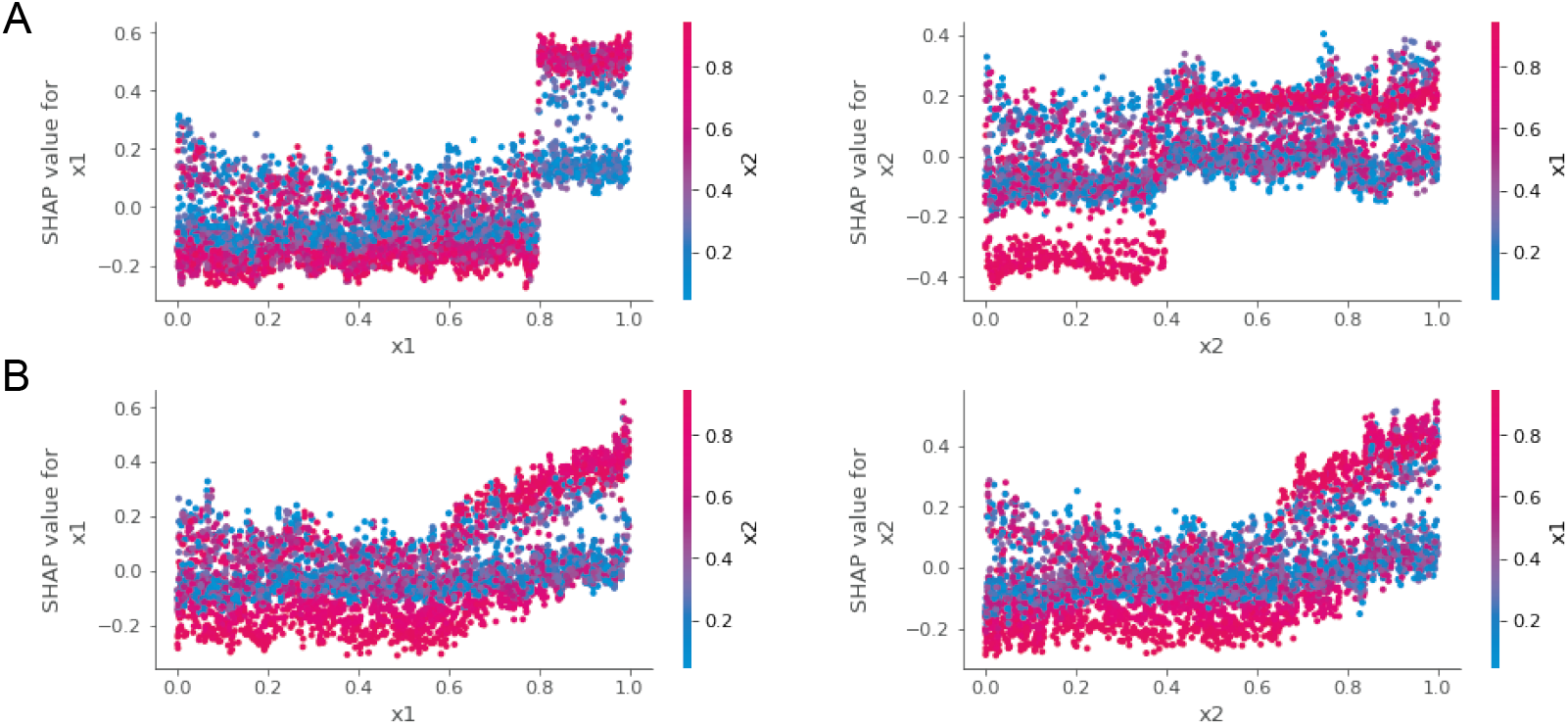
Tree SHAP values for models with noise, Bern(0.2). (A) Binary classifier with AND feature interaction. (B) Binary classifier with additive feature interaction.

## 6 Conclusion

Model agnostic approaches, such as Shapley values or SHAP, have improved our understanding of black-box models. Here we explored using simple two-feature binary classifiers on how the importance of a feature can be decomposed into its main and interaction effects as well as its marginal contributions. The collective effect, i.e. Shapley value, for a given feature can be readily examined in dependence plots with the underlying decomposed effects remaining prominent. Furthermore, specific interactions exhibit distinct behaviors, such as logical interactions with step-function like dependence or linearly additive interactions with linear dependence. ’In-region’ interactions further magnify the Shapley values. While the models here were restricted to two-feature models, the underlying decomposition, and their effects are translatable to multi-feature models. Together, we see the inter-connectivity between underlying model structure and Shapley values/interaction patterns.

## Footnotes

1 The feature independence assumption was made in the SHAP value calculations^1^. More recently, Aas et al^6^ have extended methods for SHAP calculations allowing for feature dependence.

2 We use *j* to refer to the other feature, given feature *i*, in the two-feature model

3 TreeSHAP^7^ was used. This is a computationally efficient algorithm for approximating SHAP values for tree-based models

## References

[1] S. Lundberg and S.-I. Lee, “A unified approach to interpreting model predictions,” in NIPS, 2017.

[2] L. S. Shapley, “A value for n-person games,” Contributions to the Theory of Games, vol. 2, pp. 307–317, 1953.

[3] J. H. Friedman and B. E. Popescu, “Predictive learning via rule ensembles,” The Annals of Applied Statistics, vol. 2, no. 3, pp. 916–954, 2008.

[4] B. M. Greenwell, B. C. Boehmke, and A. J. McCarthy, “A simple and effective model-based variable importance measure,” 2018.

[5] G. Hooker, “Discovering additive structure in black box functions,” in Proceedings of the Tenth ACM SIGKDD International Conference on Knowledge Discovery and Data Mining, KDD ’04, (New York, NY, USA), p. 575–580, Association for Computing Machinery, 2004.

[6] K. Aas, M. Jullum, and A. Løland, “Explaining individual predictions when features are dependent: More accurate approximations to shapley values,” 2020.

[7] S. M. Lundberg, G. G. Erion, and S.-I. Lee, “Consistent individualized feature attribution for tree ensembles,” arXiv preprint arXiv:1802.03888, 2018.

[8] K. Fujimoto, I. Kojadinovic, and J.-L. Marichal, “Axiomatic characterizations of probabilistic and cardinal-probabilistic interaction indices,” Games and Economic Behavior, vol. 55, pp. 72–99, 2006.

[9] M. Grabisch and M. Roubens, “An axiomatic approach to the concept of interaction among players in cooperative games,” Int. Journal of Game Theory, 1999.

